# Sampling SARS-CoV-2 proteomes for predicted CD8 T-cell epitopes as a tool for understanding immunogenic breadth and rational vaccine design

**DOI:** 10.1101/2020.08.15.250647

**Authors:** Jonathan Hare, David Morrison, Morten Nielsen

## Abstract

Predictive models for vaccine design have become a powerful and necessary resource for the expeditiousness design of vaccines to combat the ongoing SARS-CoV-2 global pandemic. Here we use the power of these predicted models to assess the sequence diversity of circulating SARS-CoV-2 proteomes in the context of an individual’s CD8 T-cell immune repertoire to identify potential. defined regions of immunogenicity. Using this approach of expedited and rational CD8 T-cell vaccine design, it may be possible to develop a therapeutic vaccine candidate with the potential for both global and local coverage.

## Introduction

The COVID-19 pandemic is a worldwide health emergency. The first cases were believed to have occurred earlier than December 2019, and as of the middle of September there have been more than 31,000,000 cases of SARS-CoV-2 infection have been reported worldwide and in excess of 965,000 deaths (Update, ecdc.2020). Vaccine development against SARS-CoV-2 is accelerating, and currently there are more than 130 vaccines in development with some progressing to clinical trials in man (Mullard, 2020). Of these vaccines the majority are for a prophylactic indication with the spike glycoprotein being the preferred antigen target.

The full repertoire of immune responses to COVID-19 in patients is still being evaluated but recent publications indicate a significant role for cell mediated immunity in clearing SARS-CoV-2 and conferring some level of protective immunity (Grifoni, Weiskopf, Ramirez, Mateus, et al., 2020). This mirrors observations made for other coronavirus infections, SARS (Li et al., 2008), MERS (Zhao et al., 2017) and other viral infections including Ebola (Sakabe et al., 2018) and Lassa virus (Sullivan et al., 2020). Concurrent with vaccine design and understanding the immune response in patients, significant efforts have been invested in understanding the virus phylogeny as it progresses around the globe to identify key sequence changes that may influence vaccine design. The data from this analysis appears to show that the majority of circulating mutations are neutral or deleterious (van Dorp et al., 2020), although a D614G mutation identified in the spike glycoprotein may cause an increase in susceptibility to infection (Korber et al., 2020)

Linking novel sequence data to the use of existing predictive *in silico* tools for epitope identification offers an intriguing approach that can complement existing vaccine design strategies. Previously we hypothesized that incorporating potential immune recognition information into established models may increase the likelihood of success. We have shown that within a population, although HLA sequences show high levels of polymorphism, there are conserved, over-represented alleles that can be used as representative of larger allele diversity (Buggert et al., 2012; Hare, Fiore-Gartland, et al., 2020)

Previously we have applied NetMHCpan (Nielsen & Andreatta, 2016) as a proxy to identify putative CD8 T-cell epitopes contained within the HIV transmitted founder virus (TFV) identified from the Protocol C clinical cohort of sub Saharan and East Africa. We have shown that it is possible to stratify and rank protein and/or proteome sequences for their contributions of potential T-cell epitopes (Ed McGowan et al., 2020). Here we propose to use the same approach to evaluate a subset of global circulating SARS-CoV-2 sequences and, using the predefined analysis applied to modeling HIV diversity, identify key regions within the SARS-CoV-2 proteome that could be included within a therapeutic T-cell vaccine.

## Methods

For genes from each SARS-CoV-2 virus proteome sequence all 8-11mer peptides were generated. The binding affinity of each peptide to the HLA alleles described above was predicted using NetMHCpan-4.1.

Binding predictions below the peptide conservation threshold were read into PostgreSQLR for analysis. The evaluated set of predicted binders was determined as peptides that appear in any virus proteomes, above the peptide conservation threshold of 2.1%. First the virus proteome with the largest number of unique predicted binders was identified. Next, the proteome that, when combined with the previously selected proteome gave the highest increase in coverage of all predicted peptide binders was included. This iterative process was repeated until 100% of all predicted T-cell epitopes were accounted for. For comparison, set-building was performed a second time using randomly selected proteomes instead of choosing the proteome that resulted in the greatest increase of peptide coverage. For more information go to dataspace.iavi.org

## Results

Our approach uses NetMHCpan to predict the HLA/peptide binding affinity for the selected virus strains and then identify a set of close binding peptides that are common across the strains. This algorithm was used to analyze 287 SARS-CoV-2 proteome sequences collected between and 08 January and 02 April 2020 and downloaded from NCBI SARS-CoV-2 virus database. The model parameters assessed included a minimum peptide conservation threshold of 2.1% (as previously defined ((Ed McGowan et al., 2020)) and a rank binding cut-off of 2.0. Figure 1 summarizes the output dataset which includes only those peptides that are present in 6 or more virus proteomes (epitope frequency) and have low rank binding scores although they may represent multiple HLA/peptide interactions (see Table 1 for input sample data and full model parameters).

**Table 1.** Model Parameters and proteome sources

| Parameter | Values |
| --- | --- |
| **SARS-COV-2 Proteomes** | Brazil (1), China (13), France (1), Greece (1), Iran (1), South Africa (1), Spain (10), Sweden (1), Taiwan (1), Turkey (1), USA (238), Unknown (16), Vietnam (2) |
| **Binding Threshold** | 1% |
| **HLA allele contributions** | 46 alleles (16 x HLA-A, 19 x HLA-B, 11 x HLA-C) |
| **HLA haplotype weighting** | None |
| **Rank Binding** | <2.0 |
| **Peptide Conservation threshold (%)** | 2.1% |
| **Peptide length** | 8, 9, 10 & 11mers |

**Figure 1.**
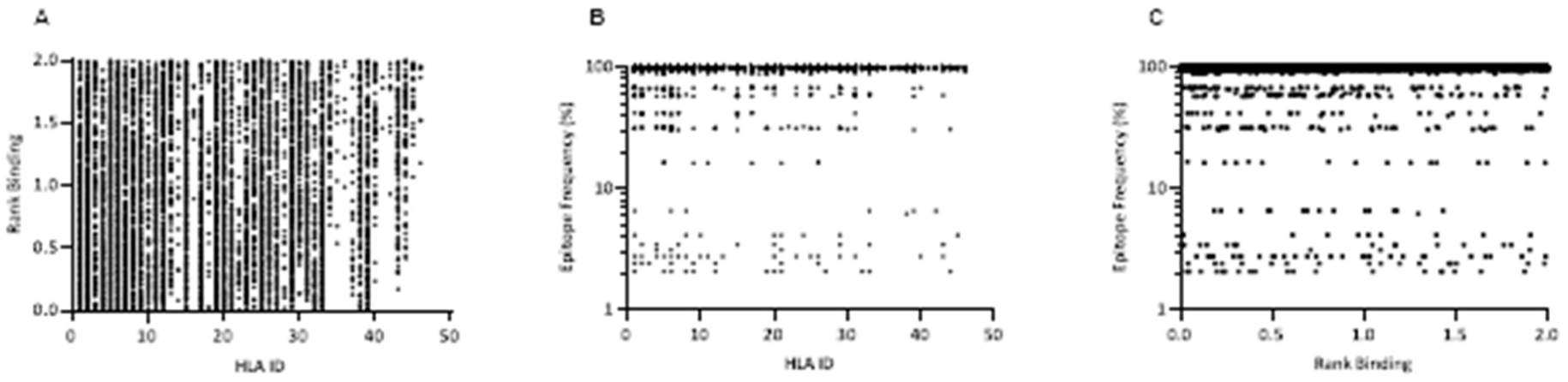
Affinity plots for all predicted peptides with conservation of ≥2.1% (n=11267). A – Predicted peptide affinity (Rank Binding) versus peptide frequency within transmitted founder proteome. B-Predicted peptide frequency versus primary associated HLA. C – Predicted peptide affinity (Rank Binding) versus primary associated HLA

These frequency and binding thresholds identified 11267 unique SARS-CoV-2-specific predicted CD8 T-cell epitopes and identified a range of predicted binding profiles for the different peptide-HLA interactions with the majority of peptides associating to selected HLA-A and HLA-B alleles (Figure 1A & B -HLA Ids 1-36) and few primary associations to HLA-C alleles.

These predicted peptides can be used to assess diversity by assigning a coverage gain value to each sequence. These values can then be used to rank each virus proteome for the coverage it provides within the sample population and by extension identify the sequences that are necessary to obtain the optimum level of epitope restricted sequence coverage (Figure 2).

**Figure 2.**
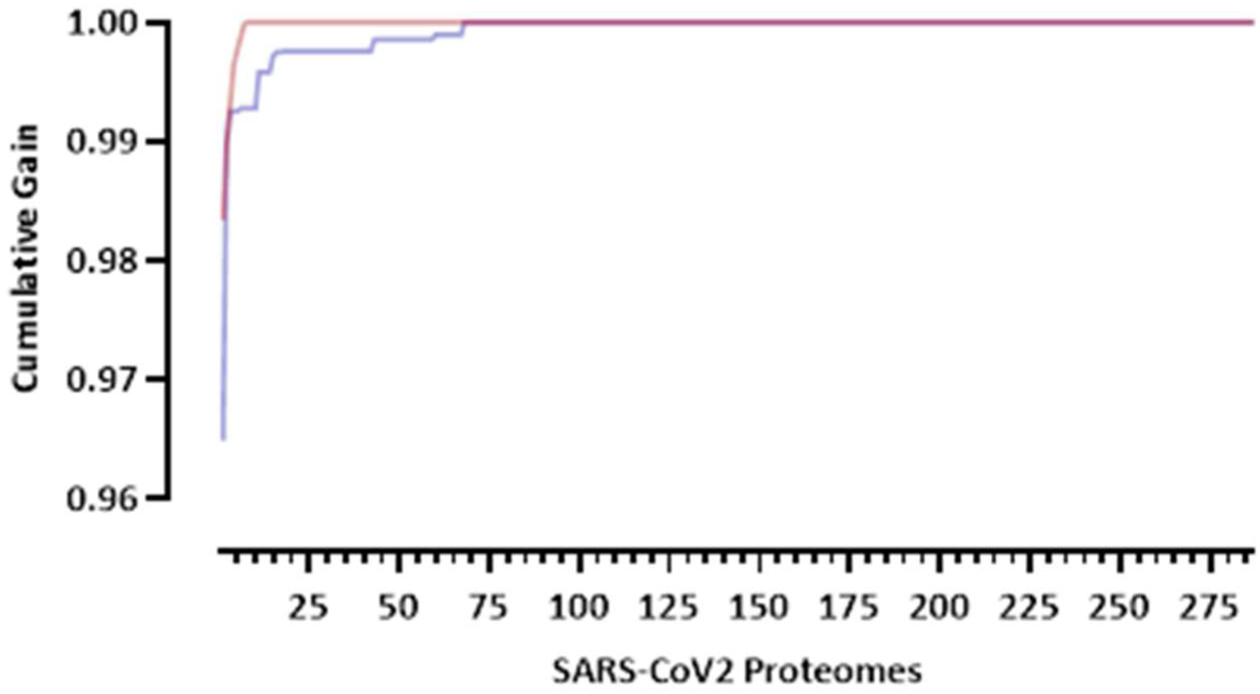
Cumulative coverage distribution plots of full length transmitted founder gag sequences using a 3-select coverage model and a 2% Binding Threshold, 3-Select best (red)) and 3-Select random (blue).

This model may be used to target and prioritize individual proteomes from which vaccine targets could be derived. In our population of 287 SARS-CoV-2 proteomes each proteome offers an individual coverage gain of 72.2-98.3%. Using this model 8 key proteomes are required to reach 100% epitope coverage within the sample population. It would require an 8-fold increase in virus sequences to achieve 100% coverage if sequences had been selected at random (n=68 p<0.0001).

Interestingly, of the 11267 unique predicted CD8 T-cell epitopes, 409 are contained in all 287 proteomes. These peptides map to 4 genes within the proteome with 392 peptides contained in ORF1ab polypeptide and spike glycoprotein (Figure 3 and Supplementary Table 1).

**Figure 3.**
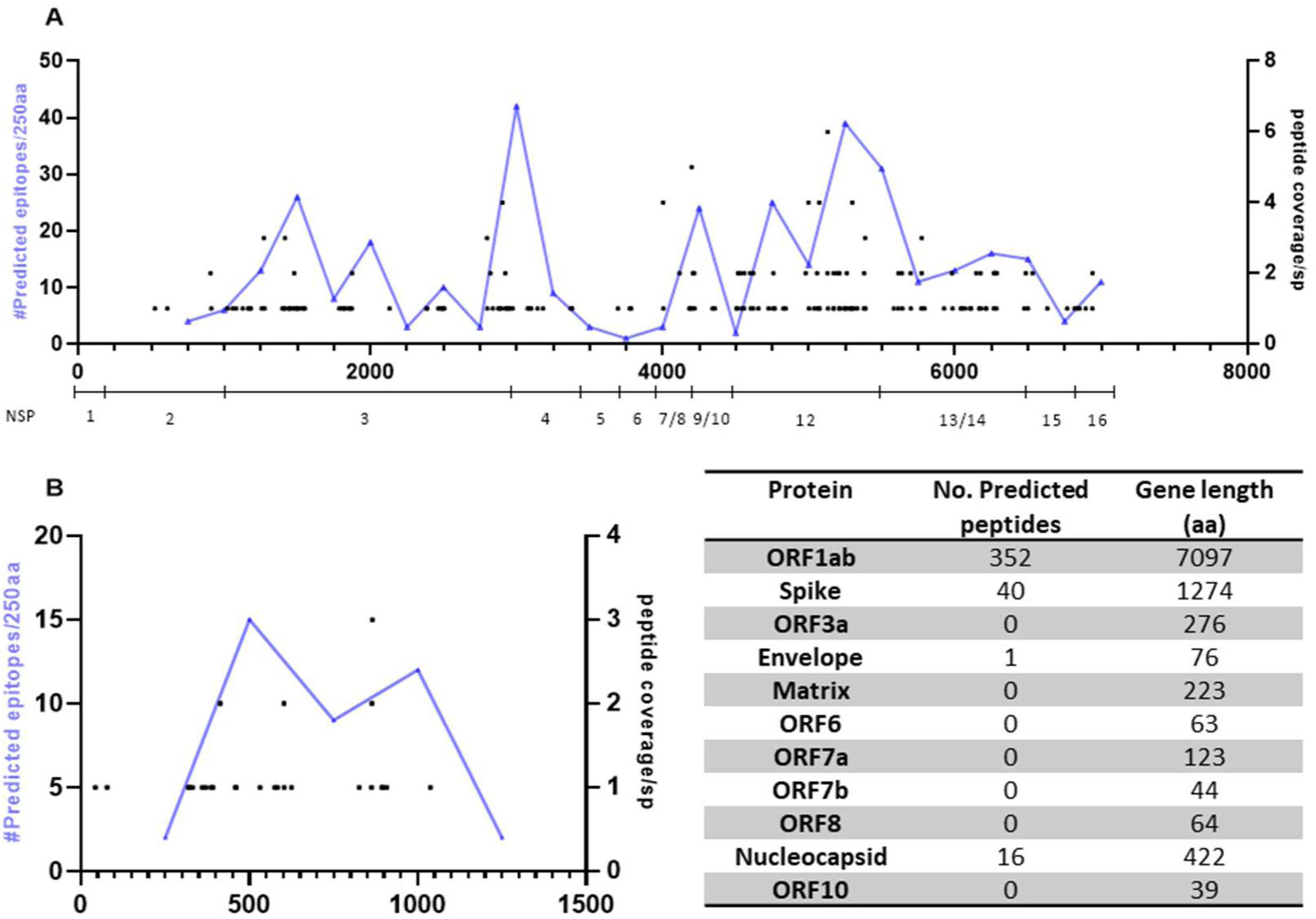
Mapping of predicted epitopes with 100% conservation showing location and number of predicted responses across the proteome and cumulative number of precited epitopes per 250aa. A – Predicted responses mapped across the ORF1ab polyprotein (with NSP genes identified). B - Predicted responses mapped across the spike glycoprotein.

## Discussion

Evaluating predicted CD8 T-cell epitopes within the SARS-CoV-2 proteome revealed a high level of conservation between proteomes, with each proteome representing 72 -98.3% of all predicted T-cell epitopes; however, using our model it is possible to reach 100% epitope coverage using 8 defined proteomes. This is in contrast to HIV where it would take 83 proteomes to achieve 100% epitope coverage on a comparable sample size (Ed McGowan et al., 2020). The enhanced conservation of epitopes is in concordance with the observed sequence diversity to date and indicates that no bias has been introduced in to the analysis through modeling with reduced input data.

Moreover, the elevated conservation can be used as a guide to identify regions within the proteome that should be included within a therapeutic T-cell vaccine. Recent data has indicated that naturally occurring T-cell responses in convalescent COVID-19 patients preferentially target the spike glycoprotein and ORF1a polypeptide (Dong et al., 2020; Grifoni, Weiskopf, Ramirez, Smith, et al., 2020). However, limitations in both these studies including that ORF1ab peptides were not examined in one study, the sample size in both was small, peptides were generated from the reference strain and the HLA Class 1 distribution was limited, may mean that potential regions of interest have been overlooked.

We identified 409 predicted CD8 peptides that have 100% conservation within our sample set with >90% of the predicted epitopes contained within either the ORF1ab polypeptide or spike glycoprotein. Furthermore, the predicted epitopes appear to cluster within a ∼550 a.a region of the spike glycoprotein (amino acid 319-865) and within 2 regions of the ORF1ab polypeptide (amino acid positions 2750-3250 and 4500-5500). Future experimental testing of these epitopes would confirm whether natural infection induces CD8 T-cell responses targeting these regions, but from an *in silico* perspective they offer a potential target for developing a therapeutic T-cell vaccine that warrants further investigation.

## Author Contributions

JH wrote the manuscript and provided conceptual input. DM and MN developed and implemented the analysis scripts and contributed to developing the manuscript.

## Acknowledgements

This work was funded in part by IAVI and made possible by the support of the United States Agency for International Development (USAID) and other donors. The full list of IAVI donors is available at http://www.iavi.org. The contents of this manuscript are the responsibility of IAVI and do not necessarily reflect the views of USAID or the US Government. We would also like to thank Thiru Thangarajah (Genscript Inc.) for insight in to potential peptide strategies. This publication was kindly accepted for pre-print publication by BioRxiv (Hare, Morrison, & Nielsen, 2020)

## References

Buggert, M., Norström, M. M., Czarnecki, C., Tupin, E., Luo, M., Gyllensten, K., … Karlsson, A. C. (2012). Characterization of HIV-specific CD4+ T cell responses against peptides selected with broad population and pathogen coverage. PLoS ONE, 7(7). https://doi.org/10.1371/journal.pone.0039874

Dong, T., Peng, Y., Mentzer, A. J., Liu, G., Yao, X., Yin, Z., … Goulder, P. (2020). Broad and strong memory CD4+ and CD8+ T cells induced by SARS-CoV-2 in UK convalescent COVID-19 patients. BioRxiv, 2020.06.05.134551. https://doi.org/10.1101/2020.06.05.134551

Grifoni, A., Weiskopf, D., Ramirez, S. I., Mateus, J., Dan, J. M., Rydyznski Moderbacher, C., … Silva, de. (2020). Journal Pre-proof Targets of T cell responses to SARS-CoV-2 coronavirus in humans with COVID-19 disease and unexposed individuals. Cell. https://doi.org/10.1016/j.cell.2020.05.015

Grifoni, A., Weiskopf, D., Ramirez, S. I., Smith, D. M., Crotty, S., & Sette, A. (2020). Targets of T Cell Responses to SARS-CoV-2 Coronavirus in Humans with COVID-19 Disease and Unexposed Individuals. https://doi.org/10.1016/j.cell.2020.05.015

Hare, J., Fiore-Gartland, A., Mcgowan, E., Rosenthal, R., Hunter, E., Gilmour, J., & Nielsen, M. (2020). Selective HLA restriction permits the evaluation and interpretation of immunogenic breadth at comparable levels to autologous HLA. https://doi.org/10.20944/preprints202008.0467.v1

Hare, J., Morrison, D., & Nielsen, M. (2020). Sampling SARS-CoV-2 proteomes for predicted CD8 T-cell epitopes as a tool for understanding immunogenic breadth and rationale 2 vaccine design. BioRxiv, 2020.08.15.250647. https://doi.org/10.1101/2020.08.15.250647

Korber, B., Fischer, W., Gnanakaran, S. G., Yoon, H., Theiler, J., Abfalterer, W., … Group, S. C.-19 G. (2020). Spike mutation pipeline reveals the emergence of a more transmissible form of SARS-CoV-2. BioRxiv, 2020.04.29.069054. https://doi.org/10.1101/2020.04.29.069054

Li, C. K., Wu, H., Yan, H., Ma, S., Wang, L., Zhang, M., … Xu, X.-N. (2008). T Cell Responses to Whole SARS Coronavirus in Humans. The Journal of Immunology, 181(8), 5490–5500. https://doi.org/10.4049/jimmunol.181.8.5490

McGowan, Ed, Rosenthal, R., Fiore-Gartland, A., Macharia, G., Balinda, S., Kapaata, A., … Hare, J. (2020). Utilizing Computational Machine Learning Tools to Understand Immunogenic Breadth in the Context of a CD8 T-Cell Mediated 2 HIV Response 3. BioRxiv, 2020.08.15.250589. https://doi.org/10.1101/2020.08.15.250589

McGowan, Edward, Rosenthal, R., Fiore-Gartland, A., Dalel, J., Streatfield, C., Coutinho, H., … Hare, J. (n.d.). Utilizing Computational Machine Learning Tools to Understand Immunogenic Breadth in the Context of a CD8 T-Cell Mediated HIV Response. Submitted.

Mullard, A. (2020). COVID-19 vaccine development pipeline gears up. https://doi.org/10.1016/S0140-6736(20)31252-6

Nielsen, M., & Andreatta, M. (2016). NetMHCpan-3.0; improved prediction of binding to MHC class I molecules integrating information from multiple receptor and peptide length datasets. Genome Medicine, 8(1), 33. https://doi.org/10.1186/s13073-016-0288-x

Sakabe, S., Sullivan, B. M., Hartnett, J. N., Robles-Sikisaka, R., Gangavarapu, K., Cubitt, B., … Oldstone, M. B. A. (2018). Analysis of CD8+ T cell response during the 2013–2016 Ebola epidemic in West Africa. Proceedings of the National Academy of Sciences of the United States of America, 115(32), E7578–E7586. https://doi.org/10.1073/pnas.1806200115

Sullivan, B. M., Sakabe, S., Hartnett, J. N., Ngo, N., Goba, A., Momoh, M., … Oldstone, M. B. A. (2020). High crossreactivity of human T cell responses between Lassa virus lineages. PLOS Pathogens, 16(3), e1008352. https://doi.org/10.1371/journal.ppat.1008352

Update, E. (n.d.). COVID-19 situation update worldwide, as of 2 August 2020. Retrieved August 3, 2020, from https://www.ecdc.europa.eu/en/geographical-distribution-2019-ncov-cases

van Dorp, L., Acman, M., Richard, D., Shaw, L. P., Ford, C. E., Ormond, L., … Balloux, F. (2020). Emergence of genomic diversity and recurrent mutations in SARS-CoV-2. Infection, Genetics and Evolution, 83, 104351. https://doi.org/10.1016/j.meegid.2020.104351

Zhao, J., Alshukairi, A. N., Baharoon, S. A., Ahmed, W. A., Bokhari, A. A., Nehdi, A. M., … Zhao, J. (2017). Recovery from the Middle East respiratory syndrome is associated with antibody and T cell responses. Science Immunology, 2(14). https://doi.org/10.1126/sciimmunol.aan5393

