## Supplementary Table 1 for "Sampling SARS-CoV-2 proteomes for predicted CD8 T-cell epitopes as a tool for understanding immunogenic breadth and rational vaccine design" (from another version of this preprint: version 2, 2020-08-17)

| **Peptide** | **Gene** | **Peptide** | **Gene** | **Peptide** | **Gene** | **Peptide** | **Gene** |
| --- | --- | --- | --- | --- | --- | --- | --- |
| AADPAMHAA | ORF1ab | GQTFSVLACY | ORF1ab | MLTNDNTSRY | ORF1ab | TNTSNQVAVLY | Glycoprotein |
| AANTVIWDY | ORF1ab | GSGVPVVDSY | ORF1ab | MLTNDNTSRYW | ORF1ab | TNVLEGSVAY | ORF1ab |
| ADQAMTQMY | ORF1ab | GSGVPVVDSYY | ORF1ab | MLVKQGDDY | ORF1ab | TPSGTWLTY | NCP |
| ADVEWKFY | ORF1ab | GSIIQFPNTY | ORF1ab | MLVKQGDDYVY | ORF1ab | TQLYLGGMSY | ORF1ab |
| AHGFELTSMKY | ORF1ab | GTGPEAGLPY | NCP | MMSAPPAQY | ORF1ab | TSNQVAVLY | Glycoprotein |
| AIDAYPLTK | ORF1ab | GTSSGDATTAY | ORF1ab | MSAPPAQY | ORF1ab | TSNQVAVLYQ | Glycoprotein |
| AIVSTIQRKY | ORF1ab | GVPVVDSYY | ORF1ab | MVMCGGSLY | ORF1ab | TSRYWEPEFY | ORF1ab |
| ALCEKALKY | ORF1ab | GVVDYGARFY | ORF1ab | NAMQVESDDY | ORF1ab | TSSGDATTAY | ORF1ab |
| ALLSDLQDLKW | ORF1ab | GVYSVIYLY | ORF1ab | NCVADYSVLY | Glycoprotein | TSSGDATTAYA | ORF1ab |
| ALLTKSSEY | ORF1ab | HADQLTPTW | Glycoprotein | NDSKEGFFTY | ORF1ab | TTAYANSVF | ORF1ab |
| AMQVESDDY | ORF1ab | HFAIGLALY | ORF1ab | NLEEAARY | ORF1ab | TTCCSLSHRFY | ORF1ab |
| ANDPVGFTL | ORF1ab | HFAIGLALYY | ORF1ab | NLQSNHDLY | ORF1ab | TTDAVRDPQTL | Glycoprotein |
| ASDTYACWH | ORF1ab | HGFELTSMKY | ORF1ab | NLVAVPTGY | ORF1ab | TTFDSEYCRH | ORF1ab |
| ASDTYACWHH | ORF1ab | HISRQRLTKY | ORF1ab | NQDLNGNWY | ORF1ab | TTIKPVTY | ORF1ab |
| ASEYTGNY | ORF1ab | HLDGEVITF | ORF1ab | NSASFSTFKCY | Glycoprotein | TTLPVNVAF | ORF1ab |
| ASFSTFKCY | Glycoprotein | HTDFSSEI | ORF1ab | NSVPWDTIANY | ORF1ab | TTNGDFLHF | ORF1ab |
| ASFYYVWKSY | ORF1ab | HTDFSSEII | ORF1ab | NTDFSRVSA | ORF1ab | TTTIKPVTY | ORF1ab |
| ASGKPVPYCY | ORF1ab | HTDFSSEIIGY | ORF1ab | NTLQCIMLVY | ORF1ab | TVKNGSIHLY | ORF1ab |
| ASIKNFKSVLY | ORF1ab | HTMLVKQGDDY | ORF1ab | NTLTLAVPY | ORF1ab | TVVIGTSKFY | ORF1ab |
| AVRDPQTLEI | Glycoprotein | IAANTVIWDY | ORF1ab | NTSNQVAVLY | Glycoprotein | VADYSVLY | Glycoprotein |
| AYESLRPDTRY | ORF1ab | ICISTKHFY | ORF1ab | NTSNQVAVLYQ | Glycoprotein | VAKYTQLCQY | ORF1ab |
| AYKIEELFYSY | ORF1ab | ICYTPSKLIEY | ORF1ab | NTSRYWEPEFY | ORF1ab | VCVDTVRTNVY | ORF1ab |
| CASEYTGNY | ORF1ab | IETISLAGSY | ORF1ab | NVAKYTQLCQY | ORF1ab | VDALCEKALKY | ORF1ab |
| CDVTDVTQLY | ORF1ab | IGVTQNVLY | Glycoprotein | NVLEGSVAY | ORF1ab | VDTDFVNEFY | ORF1ab |
| CTCGKQATKY | ORF1ab | IICISTKHFY | ORF1ab | PINPTDQSSY | ORF1ab | VDTDFVNEFYA | ORF1ab |
| CTFLLNKEMY | ORF1ab | ILDITPCSF | Glycoprotein | PKSDGTGTIY | ORF1ab | VDTVRTNVY | ORF1ab |
| CVADYSVLY | Glycoprotein | ILLNKHIDAY | NCP | PKSDGTGTIYT | ORF1ab | VDTVSALVY | ORF1ab |
| CVDTVRTNVY | ORF1ab | ILMTARTVY | ORF1ab | PLLTDEMIAQY | Glycoprotein | VDYGARFY | ORF1ab |
| CYTPSKLIEY | ORF1ab | INPTDQSSY | ORF1ab | PLTKHPNQEY | ORF1ab | VENPDILRVY | ORF1ab |
| DALCEKALKY | ORF1ab | IPFAMQMAY | Glycoprotein | PQADVEWKFY | ORF1ab | VENPHLMGWDY | ORF1ab |
| DAQPCSDKAY | ORF1ab | ISDEVARDL | ORF1ab | PTDQSSYIV | ORF1ab | VFDEISMATNY | ORF1ab |
| DASGKPVPY | ORF1ab | ISDEVARDLSL | ORF1ab | QADVEWKFY | ORF1ab | VFSAVGNICY | ORF1ab |
| DASGKPVPYCY | ORF1ab | ISDYDYYRY | ORF1ab | QAENVTGLF | ORF1ab | VKNGSIHLY | ORF1ab |
| DAVTAYNGY | ORF1ab | ISDYDYYRYN | ORF1ab | QGDDYVYLPY | ORF1ab | VLEGSVAY | ORF1ab |
| DEISMATNY | ORF1ab | ISMMGFKMNY | ORF1ab | QIPFAMQMAY | Glycoprotein | VLMDGSIIQF | ORF1ab |
| DELTGHMLDMY | ORF1ab | ISRQRLTKY | ORF1ab | QLTPTWRVY | Glycoprotein | VLPFNDGVY | Glycoprotein |
| DIQLLKSAY | ORF1ab | ITDVFYKENSY | ORF1ab | QLYLGGMSY | ORF1ab | VMLTNDNTSRY | ORF1ab |
| DLSPRWYFY | NCP | ITFDNLKTLL | ORF1ab | QLYLGGMSYY | ORF1ab | VMMSAPPAQY | ORF1ab |
| DLSPRWYFYY | NCP | ITFLKKDAPY | ORF1ab | QTFSVLACY | ORF1ab | VPFVVSTGY | ORF1ab |
| DNQDLNGNWY | ORF1ab | IVDTVSALV | ORF1ab | QTGKIADY | Glycoprotein | VPQADVEWKFY | ORF1ab |
| DQFKHLIPLMY | ORF1ab | IVDTVSALVY | ORF1ab | QTGKIADYNY | Glycoprotein | VPWDTIANY | ORF1ab |
| DQLTPTWRVY | Glycoprotein | IVDTVSALVYD | ORF1ab | QTIEVNSFSGY | ORF1ab | VSDIDITF | ORF1ab |
| DSKEGFFTY | ORF1ab | IVSTIQRKY | ORF1ab | QWLTNIFGTVY | ORF1ab | VSDIDITFL | ORF1ab |
| DTDFVNEF | ORF1ab | KAYKIEELFY | ORF1ab | RFDNPVLPF | Glycoprotein | VSDIDITFLK | ORF1ab |
| DTDFVNEFY | ORF1ab | KDASGKPVPY | ORF1ab | RGDKSVYY | ORF1ab | VSEETGTLI | Envelope |
| DTDFVNEFYA | ORF1ab | KDLSPRWYFY | NCP | RLSFKELLVY | ORF1ab | VSIINNTVY | ORF1ab |
| DTDFVNEFYAY | ORF1ab | KIEELFYSY | ORF1ab | RPDTRYVLM | ORF1ab | VSSPDAVTAY | ORF1ab |
| DTLKEILVTY | ORF1ab | KIQEGVVDY | ORF1ab | RPINPTDQSSY | ORF1ab | VTDTPKGPK | ORF1ab |
| DTNVLEGSVAY | ORF1ab | KLNDLCFTNVY | Glycoprotein | RVDFCGKGY | Glycoprotein | VTDTPKGPKV | ORF1ab |
| DTPKGPKVKY | ORF1ab | KMADQAMTQMY | ORF1ab | RVDWTIEY | ORF1ab | VTDVTQLY | ORF1ab |
| DTVRTNVY | ORF1ab | KMKDLSPRWY | NCP | RVFSAVGNICY | ORF1ab | VTDVTQLYL | ORF1ab |
| DTVSALVY | ORF1ab | KMNYQVNGY | ORF1ab | RVVTTFDSEY | ORF1ab | VTPSGTWLTY | NCP |
| DVDTDFVNEF | ORF1ab | KNFKSVLYY | ORF1ab | SAQCFKMFY | ORF1ab | VTQLYLGGMSY | ORF1ab |
| DVDTDFVNEFY | ORF1ab | KQGDDYVYLPY | ORF1ab | SASFSTFKCY | Glycoprotein | VTTFDSEY | ORF1ab |
| DVFYKENSY | ORF1ab | KRVDFCGKGY | Glycoprotein | SDGTGTIY | ORF1ab | VTVKNGSIHLY | ORF1ab |
| DVTDVTQLY | ORF1ab | KSAQCFKMFY | ORF1ab | SDYDYYRY | ORF1ab | VTYNCCDDDY | ORF1ab |
| DVTDVTQLYL | ORF1ab | KSDGTGTIY | ORF1ab | SEMVMCGGSLY | ORF1ab | VVAFNTLLF | ORF1ab |
| EFKLASHMY | ORF1ab | KSDGTGTIYT | ORF1ab | SGDATTAY | ORF1ab | VVDSYYSLL | ORF1ab |
| EGVVDYGARFY | ORF1ab | KSDGTGTIYTE | ORF1ab | SGEFKLASHMY | ORF1ab | VVDSYYSLLM | ORF1ab |
| EIVDTVSALVY | ORF1ab | KSHFAIGLALY | ORF1ab | SGKPVPYCY | ORF1ab | VVDYGARFY | ORF1ab |
| ELIRQGTDY | NCP | KTDGTLMIERF | ORF1ab | SGVPVVDSY | ORF1ab | VVDYGARFYF | ORF1ab |
| ELTGHMLDMY | ORF1ab | KVDGVDVEL | ORF1ab | SGVPVVDSYY | ORF1ab | VVDYGARFYFY | ORF1ab |
| EMVMCGGSLY | ORF1ab | KVDGVDVELF | ORF1ab | SIIQFPNTY | ORF1ab | VVIGTSKFY | ORF1ab |
| ENDSKEGFFTY | ORF1ab | KVVKVTIDY | ORF1ab | SIKNFKSVLY | ORF1ab | VVTTFDSEY | ORF1ab |
| ENPDILRVY | ORF1ab | LAVFDKNLY | ORF1ab | SIKNFKSVLYY | ORF1ab | VVVNAANVY | ORF1ab |
| EPEFYEAMY | ORF1ab | LDNQDLNGNWY | ORF1ab | SKRVDFCGKGY | Glycoprotein | VYSDVENPHLM | ORF1ab |
| EQKSILSPLY | ORF1ab | LLEKCDLQNY | ORF1ab | SMDNSPNLA | ORF1ab | WFSQRGGSY | ORF1ab |
| ESLRPDTRY | ORF1ab | LLNKHIDAY | NCP | SMDNSPNLAW | ORF1ab | WLTNIFGTVY | ORF1ab |
| ETDLTKGPH | ORF1ab | LLSDLQDLKW | ORF1ab | SMMGFKMNY | ORF1ab | YDAQPCSDKAY | ORF1ab |
| ETDLTKGPHEF | ORF1ab | LLTDEMIAQY | Glycoprotein | SNCVADYSVLY | Glycoprotein | YESLRPDTRY | ORF1ab |
| ETISLAGSY | ORF1ab | LLTDEMIAQYT | Glycoprotein | SNQVAVLY | Glycoprotein | YFDKAGQKTY | ORF1ab |
| ETLVTMPLGY | ORF1ab | LMDGSIIQF | ORF1ab | SPDAVTAY | ORF1ab | YIDINGNLH | ORF1ab |
| EVTPSGTWLTY | NCP | LNDLCFTNVY | Glycoprotein | SPDAVTAYNGY | ORF1ab | YIFFASFYY | ORF1ab |
| FAIGLALYY | ORF1ab | LNLEEAARY | ORF1ab | SRLSFKELLVY | ORF1ab | YIICISTKHFY | ORF1ab |
| FCDLKGKY | ORF1ab | LQDLKWARF | ORF1ab | SSGDATTAY | ORF1ab | YIKWDLLKY | ORF1ab |
| FDEISMATNY | ORF1ab | LSDLQDLKW | ORF1ab | SSGDATTAYA | ORF1ab | YKIEELFYSY | ORF1ab |
| FDKAGQKTY | ORF1ab | LSDRVVFVL | ORF1ab | SSPDAVTAY | ORF1ab | YLAVFDKNLY | ORF1ab |
| FHLDGEVITF | ORF1ab | LSFKELLVY | ORF1ab | SSSKTPEEH | ORF1ab | YLFDESGEF | ORF1ab |
| FIETISLAGSY | ORF1ab | LSPRWYFY | NCP | STDTCFANK | ORF1ab | YLFDESGEFK | ORF1ab |
| FKMNYQVNGY | ORF1ab | LSPRWYFYY | NCP | STDTCFANKH | ORF1ab | YLGGMSYY | ORF1ab |
| FLGYFCTCY | ORF1ab | LTDEMIAQY | Glycoprotein | SVELKHFFF | ORF1ab | YLNTLTLAVPY | ORF1ab |
| FLKRGDKSVYY | ORF1ab | LTDEMIAQYT | Glycoprotein | SVPWDTIANY | ORF1ab | YNGSPSGVY | ORF1ab |
| FLPGVYSVIY | ORF1ab | LTDEMIAQYTS | Glycoprotein | SVSSPDAVTAY | ORF1ab | YPLTKHPNQEY | ORF1ab |
| FPKSDGTGTIY | ORF1ab | LTGHMLDMY | ORF1ab | SYTTTIKPVTY | ORF1ab | YSDVENPHL | ORF1ab |
| FSAVGNICY | ORF1ab | LTKHPNQEY | ORF1ab | TCCSLSHRFY | ORF1ab | YSDVENPHLM | ORF1ab |
| FSQRGGSY | ORF1ab | LTKYTMADLVY | ORF1ab | TCGKQATKY | ORF1ab | YSDVENPHLMG | ORF1ab |
| FSSEIIGY | ORF1ab | LTNDNTSRY | ORF1ab | TDEMIAQY | Glycoprotein | YSFLPGVY | ORF1ab |
| FTNVYADSF | Glycoprotein | LTNDNTSRYW | ORF1ab | TDFSSEIIGY | ORF1ab | YTDFATSA | ORF1ab |
| FTVLCLTPVY | ORF1ab | LTNIFGTVY | ORF1ab | TDFVNEFY | ORF1ab | YTDFATSAC | ORF1ab |
| FVDGVPFVV | ORF1ab | LVDSDLNDF | ORF1ab | TDFVNEFYAY | ORF1ab | YTDFATSACV | ORF1ab |
| FVENPDILRVY | ORF1ab | LVKQGDDYVY | ORF1ab | TDTPKGPKVKY | ORF1ab | YTDFATSACVL | ORF1ab |
| FVKRVDWTIEY | ORF1ab | LYDKLVSSF | ORF1ab | TDVFYKENSY | ORF1ab | YTELEPPCR | ORF1ab |
| FVMMSAPPAQY | ORF1ab | LYFDKAGQKTY | ORF1ab | TDVTQLYL | ORF1ab | YTELEPPCRF | ORF1ab |
| FVNEFYAY | ORF1ab | MADLVYALR | ORF1ab | TGNLQSNHDLY | ORF1ab | YTGNYQCGHY | ORF1ab |
| FVSLAIDAY | ORF1ab | MADLVYALRH | ORF1ab | TGNYQCGHY | ORF1ab | YTKVDGVDVEL | ORF1ab |
| GCDVTDVTQLY | ORF1ab | MADQAMTQM | ORF1ab | TGPEAGLPY | NCP | YTMADLVY | ORF1ab |
| GDDYVYLPY | ORF1ab | MADQAMTQMY | ORF1ab | TIEVNSFSGY | ORF1ab | YTPSKLIEY | ORF1ab |
| GEFKLASHMY | ORF1ab | MADQAMTQMYK | ORF1ab | TLKEILVTY | ORF1ab | YTTTIKPVTY | ORF1ab |
| GFELTSMKY | ORF1ab | MIDVQQWGF | ORF1ab | TLQCIMLVY | ORF1ab | YVDTPNNTDF | ORF1ab |
| GIGVTQNVLY | Glycoprotein | MKDLSPRWY | NCP | TLVTMPLGY | ORF1ab | YVLMDGSIIQF | ORF1ab |
| GIMMNVAKY | ORF1ab | MKDLSPRWYFY | NCP | TMLVKQGDDY | ORF1ab | YWEPEFYEAMY | ORF1ab |
| GLNLEEAARY | ORF1ab | MLDMYSVML | ORF1ab | TNDNTSRY | ORF1ab |  |  |
| GNLQSNHDLY | ORF1ab | MLLEKCDLQNY | ORF1ab |  |  |  |  |

Supplementary Table 1. Predicted peptides and corresponding genes with 100% prevalence in analyzed cohort
